# Mapping the Cell-Membrane Proteome to the Cancer Hallmarks

**DOI:** 10.1101/2022.03.18.484818

**Authors:** Iulia M. Lazar, Arba Karcini, Joshua R. S. Haueis

## Abstract

The hallmarks of biological processes that underlie the development of cancer have been long recognized, yet, existing therapeutic treatments cannot prevent cancer from continuing to be one of the leading causes of death worldwide. This work was aimed at exploring the extent to which the cell-membrane proteins are implicated in triggering cancer hallmark processes, and assessing the ability to pinpoint novel therapeutic targets through a combined membrane proteome/cancer hallmark perspective. By using GO annotations, a database of human proteins associated broadly with ten cancer hallmarks was created. Cell-membrane cellular subfractions of SKBR3/HER2+ breast cancer cells used as a model system were analyzed by high resolution mass spectrometry, and high-quality proteins (FDR<3 %) identified by at least two unique peptides were mapped to the cancer hallmark database. Over 1100 experimentally detected cell-membrane or cell-membrane associated proteins, representing ~8 % of the human cell-membrane proteome, were mapped to the hallmark database. Representative membrane constituents such as receptors, CDs, adhesion and transport proteins were distributed over the entire genome and present in every hallmark category. Sustained proliferative signaling/cell cycle, adhesion/tissue invasion, and evasion of immune destruction emerged as prevalent hallmarks represented by the membrane proteins. Construction of protein-protein interaction networks uncovered a high level of connectivity between the hallmark members, with some receptor (EGFR, ERBB2, FGFR, MTOR, CSF1R), antigen (CD44), and adhesion (MUC1) proteins being implicated in most hallmark categories. An illustrative subset of 116 hallmark proteins that included 44 oncogenes, 28 tumor suppressors, and 38 approved drug targets was subjected to a more in-depth analysis. The existing drug targets were implicated mainly in signaling processes. Network centrality analysis revealed that nodes with high degree, rather than betweenness, represent a good resource for informing the selection of putative novel drug targets. Through heavy involvement in supporting cancer hallmark processes, we show that the functionally diverse and networked landscape of cancer cell-membrane proteins fosters unique opportunities for guiding the development of novel therapeutic interventions, including multi-agent, immuno-oncology and precision medicine applications.

## Introduction

The cancer hallmarks were first described by Hanahan and Weinberg [1,2], and evolved to include six fundamental biological capabilities that sustain neoplastic growth (sustained proliferative signaling, insensitivity to anti-growth signals, evasion of apoptosis, limitless replicative potential, sustained angiogenesis, tissue invasion and metastasis), two emerging hallmarks of a more general nature (reprogramming of energy metabolism and evasion of immune destruction), and two enabling characteristics (genome instability and inflammation). Genetic changes drive the formation of a primary tumor, however, cell-autonomous mechanisms propelled by a diverse mutational landscape are not sufficient to explain the full range of aberrant behaviors. For example, tissue invasion and metastatic propensity have been described as being driven by a supportive tumor micro- or systemic macro-environment and epigenetic re-programming rather than metastasis-specific driver mutations [3–7]. Mutations, on the other hand, have been shown to heavily affect epigenetic regulators [8]. Also, synergistic effects enabled by cell-extrinsic environmental stimuli (various signaling, growth, cytokine or angiogenic factors, recruited stromal cells, etc.) and cell-intrinsic EMT gene regulators have been proposed to drive the metastatic processes via adaptation rather than selection [8–11]. As a result, in support of the metastatic process, additional hallmarks that include microenvironment modulation, plasticity, motility/invasion, and colonization have been proposed [12]. Moreover, the presence of a tumor microbiome has been implicated in tumor-supportive inflammation and tumor progression [13]. Altogether, genetic and epigenetic alterations work in tandem with a cooperative environment to reprogram gene expression, corrupt biological regulatory pathways, mediate adaptation in support of malignant neoplastic growth and metastasis, and ultimately drive the evolution of drug resistance [8,14].

Non-cell-autonomous mechanisms that support cancer development and steer its evolution act through the interface between the cancer cells and their environment. This interface is defined by the cell-membrane that imbeds proteins with critical roles in intra-cellular signaling and inter-cellular communication, cell-cell, cell-pathogen and immune recognition events, cell-adhesion and motility, and exchange of solutes and various factors. Cell-surface receptor tyrosine kinases (RTKs) trigger vital signaling cascades that control proliferation, differentiation, growth and metabolism. RTKs initiate the signaling process and regulate intracellular events by binding various ligands such as growth factors, peptides and hormones. Aberrant or mutated expression of such receptors is often the driver of uncontrolled proliferation. In contrast, cytokine receptors initiate signaling processes through association with other non-receptor protein kinases, while membrane proteases support secretory, cell-cell signaling, and degradation functions by cleaving the membrane-bound proteins. Plasma membrane proteins also interact with lipids and carbohydrates to sustain various transport and endo-/exocytic processes to ensure in-and-out shuffling of solutes, nutrients, hormones, growth factors, cytokines, and other signaling and ECM molecules. Cell-junction and adhesion proteins, on the other hand, are critical not just to determining the 3D architecture of cell conglomerates, but also to inter-cellular or cell-ECM communication, as well as functionality within a tissue. As a result, in the context of disease, the study of an extensive catalogue of cell-membrane proteins through their collective involvement in disease mechanisms, rather than specific functional roles, would be more meaningful to developing effective cures.

In this work, SKBR3/HER2+ cells were used as a model system to explore the cell-membrane proteome through the perspective of cancer hallmarks. Cell-membrane proteomic data generated by complementary methods were analyzed and queried for the presence of proteins that could be correlated with the hallmark processes. The results were assessed in the context of emerging interest in novel therapeutic paradigms that seek a systems approach for identifying drug targets or drug target combinations with synergistic effects. Protein-protein interaction (PPI) network centrality measures indicated the presence of potentially novel and valuable cancer drug targets.

## Methods

### Cell-membrane protein fraction preparation and analysis

A detailed description of the experimental conditions used for cell culture, cell-membrane protein labeling/isolation, mass spectrometry (MS) analysis, and annotation of membrane proteins is provided in reference [15]. Briefly, SKBR3 breast cancer cells were acquired from ATCC (Manassas, VA), authenticated by STR (ATCC), grown either for 48 h in serum-free (McCoy 5A, Gibco, Carlsbad, CA) or 48 h serum-free/24 h serum-rich (10 % FBS, Gemini Bioproducts, West Sacramento, CA) culture media at 37° C/5 % CO_2_, and processed for generating cell-membrane enriched protein fractions. The SKBR3 cell-membrane proteins were isolated by either the biotinylation of amino (EZ-Link Sulfo-NHS-SS-Biotin, Thermo Fisher Scientific, Rockford, IL) or oxidized glycan (EZ-Link Alkoxamine-PEG4-Biotin, Thermo Fisher Scientific) groups followed by affinity NeutrAvidin (Thermo Fisher Scientific) pulldown, or by tryptic shaving of cell-surface proteins by using recombinant enzymes (TrypLE, Gibco). Three biological replicates of cells cultured in the presence or absence of serum, enriched in membrane proteins according to all three protocols, were generated. The tryptic proteolytic digests of the isolated proteins from all biological cell states and replicates were analyzed in triplicate by mass spectrometry using an EASY-nano liquid chromatography (LC) 1200 UHPLC/QE-Orbitrap-MS system (Thermo Fisher Scientific) and data-dependent higher energy collision (HCD)/MS2 data acquisition. The mass spectrometry raw files were deposited in the PRIDE Archive (see below access info) and together with the associated protein lists [15] were used in this study to map the detectable membrane proteins in the SKBR3 breast cancer cell line to the cancer hallmark processes.

### MS data processing

The LC-MS/MS raw files were analyzed by using the ProteomeDiscoverer 2.4 software package (Thermo Fisher Scientific) [15], the Sequest HT search engine, and a reviewed, minimally redundant *Homo sapiens* database from UniProt with 20,433 entries (2019) [16]. The MS searches were enabled for a parent peptide ion mass range of 400-5000 Da, allowing for two missed tryptic cleavages, Met oxidation, and Nt acetylation. The quality of protein IDs that were mapped to the hallmark supportive database was determined by: (a) the stringency of peptide identifications as defined by parent/fragment ion mass tolerances of 15 ppm and 0.02 Da, respectively; (b) the use of only rank 1 peptides and top scoring proteins; and (c) the use of a minimum number of two unique peptide matches to a protein sequence. All FDR targets for peptide spectrum matches, peptide and protein groups were set to either 0.03 (relaxed) or 0.01 (strict), and protein grouping was accomplished by using the strict parsimony principle. The results of all raw file database searches were combined to increase the number and confidence of protein identifications in the cell membrane. This strategy led to the identification of a dataset of 1316 protein matched by two unique peptides. Protein abundance was assessed based on the summed peptide spectrum counts (SC), as log transformed values, i.e., log10 (SC). Protein assignments to the cell-membrane or cell surface were made by using UniProt/GO annotations, the Human Protein Atlas cellular and organelle proteome database, and literature reports [15–18].

### Bioinformatics data processing

The database defining the biological processes that support the cancer hallmarks was constructed by retrieving the relevant processes and pathways from the UniProt database by using the advanced search tool with the following options: (a) *Homo sapiens* organism, (b) only reviewed UniProt entries (i.e., Swiss-Prot entries), (c) GO annotation controlled vocabulary terms, and (d) any assertion method [16,17]. The protein entries associated with these processes were extracted in the period July 2020-June 2021. Specific GO term definitions assigned to a particular cancer hallmark are provided in the table. The COSMIC (v94) Cancer Gene Census catalogue (CGC) [19], along with other hallmark proteins reported in the literature [1, 2, 20–24], were used for generating a better curated definition of hallmark proteins. Experimentally measured cell-membrane proteins were mapped to the cancer hallmark processes by aligning the list of detectable SKBR3 membrane proteins with the protein entries from the hallmark database (**Supplemental file 1**). The circular data plot representing the detected SKBR3 cell-membrane proteins mapped to the cancer hallmarks and the corresponding genes within the 23 chromosomes in the human genome was created using the Galaxy platform [25] and the Circos data visualizing package [26]. Gene start and gene end positions were determined based on Ensemble gene annotations. Protein-protein interaction networks were created in STRING [27]. All interactions sources were enabled and the minimum required interaction scores were set to high confidence (i.e., ≥0.7). Network analysis was performed with the Cytoscape NetworkAnalyzer plugin. Network visualization and attribute circle layouts based on degree and betweenness centrality measures were created with Cytoscape 3.9.1 tools [28]. This work did not involve the use of human subjects, and did not require IRB approval.

## Results

Given the critical role of the plasma membrane in determining the fate of a cell, we hypothesized that many proteins that are integral to- or associated with the plasma membrane can be re-evaluated in the context of cancer hallmarks to support the detection and therapeutic treatment of cancerous cell states. To this end, an in-house database (DB) comprising ten broad categories of proteins that are either actively participating in-, or are just supportive or enablers of the cancer hallmarks was created by using the UniProt *Homo sapiens* DB and GO biological process and pathway annotations. **Table 1** provides the original hallmark categories, the biological processes and pathways associated with the specific hallmarks, and the number of proteins matched to each category. Overall, 6,258 cell-membrane/cell-surface proteins were associated with the hallmark DB (**Figure 1A** and **Supplemental file 1**). The number of experimentally detected cell-membrane proteins, and examples of cell-membrane hallmark proteins described in the literature [20–24] or reported in the CGC DB with cancer promoting, suppressing or dual role [19], are also provided. Decision for associating a particular biological process or pathway with a hallmark was made based on relevance and availability of adequate GO annotations, with the assumption that GO annotations are accurate.

**Table 1.**
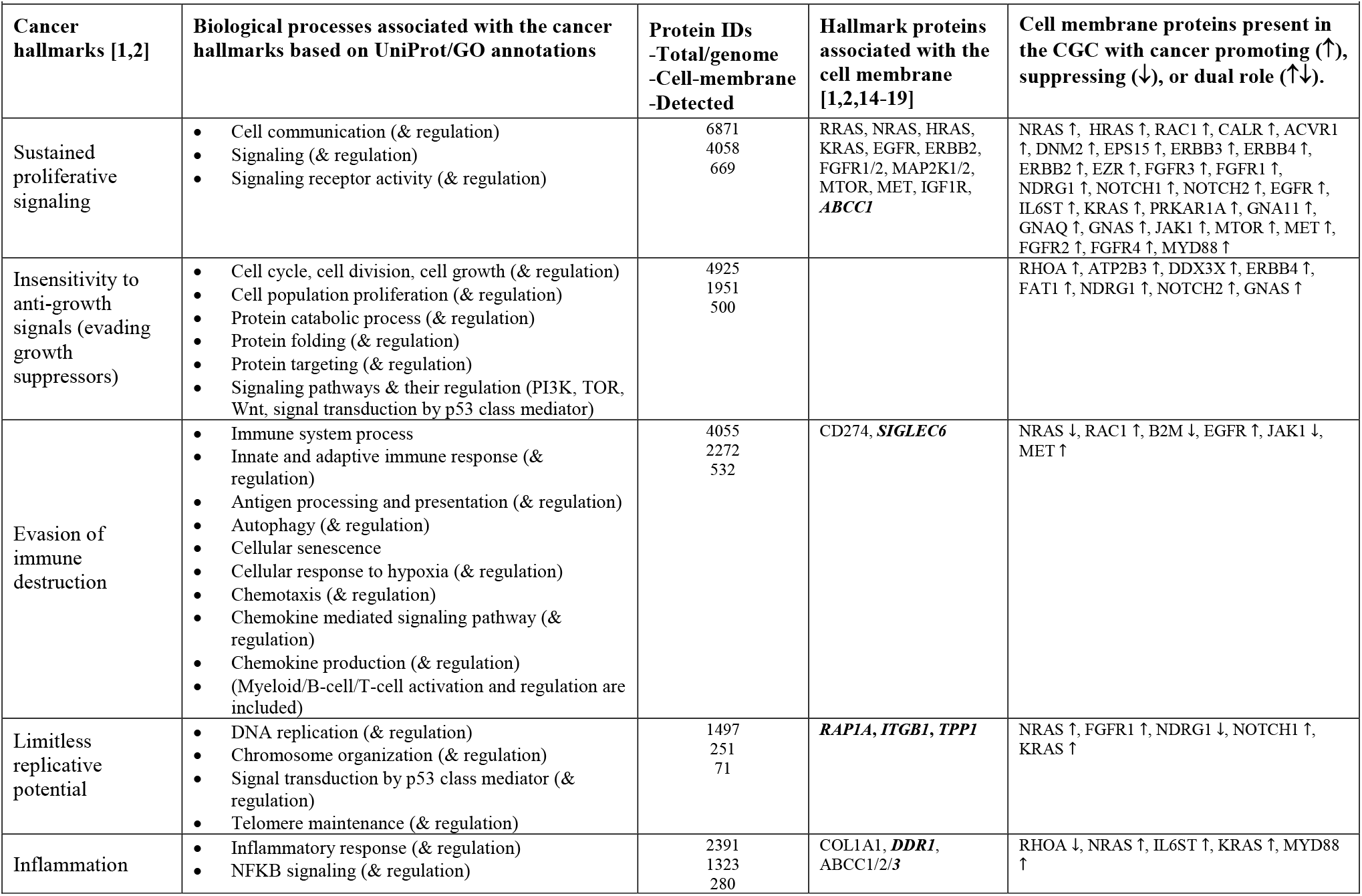
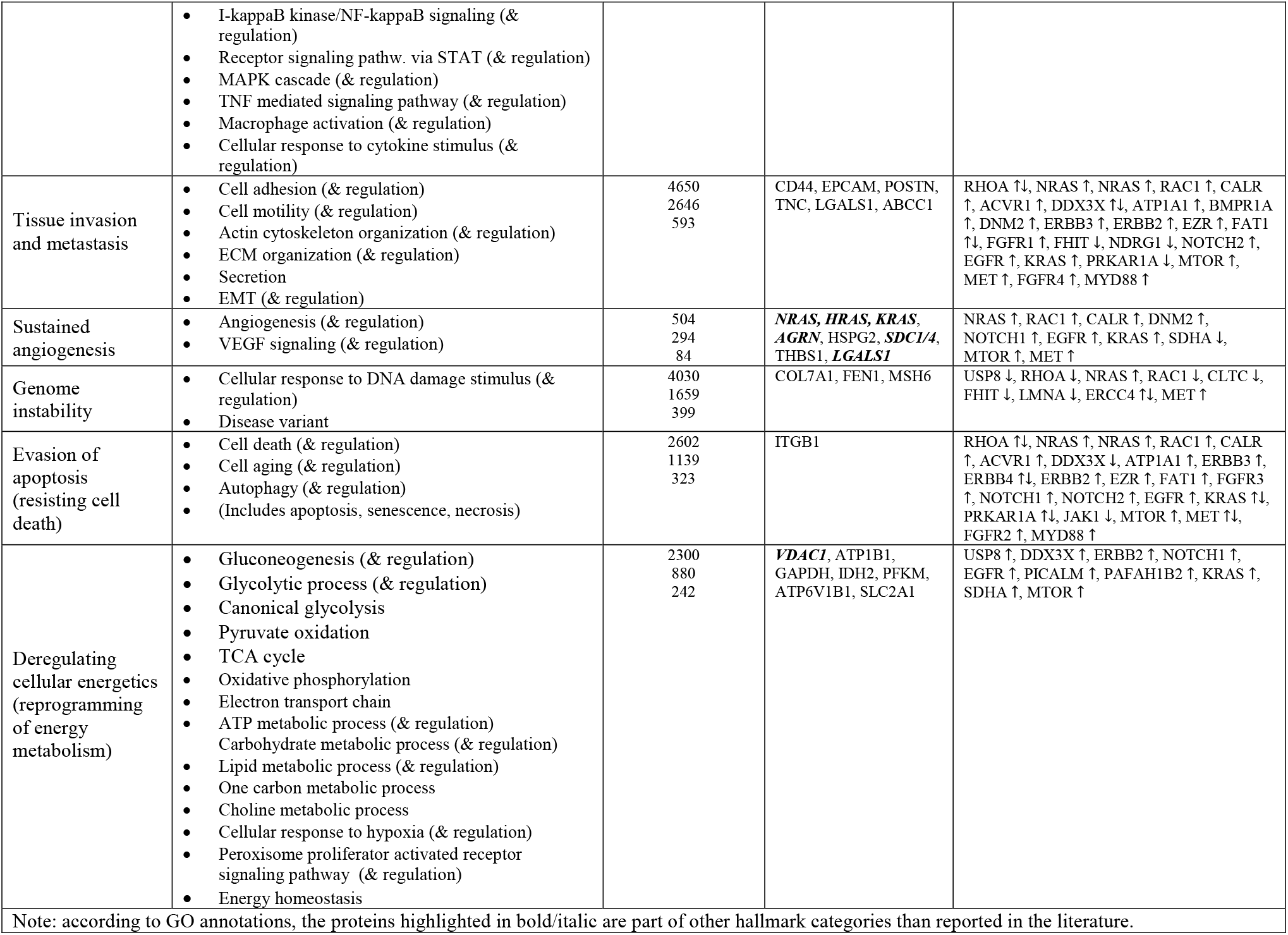
Cancer hallmarks defined by GO biological processes and examples of cell-membrane proteins associated with the hallmarks. Proteins catalogued in the CGC have promoting, suppressing, or dual role.

**Figure 1.**
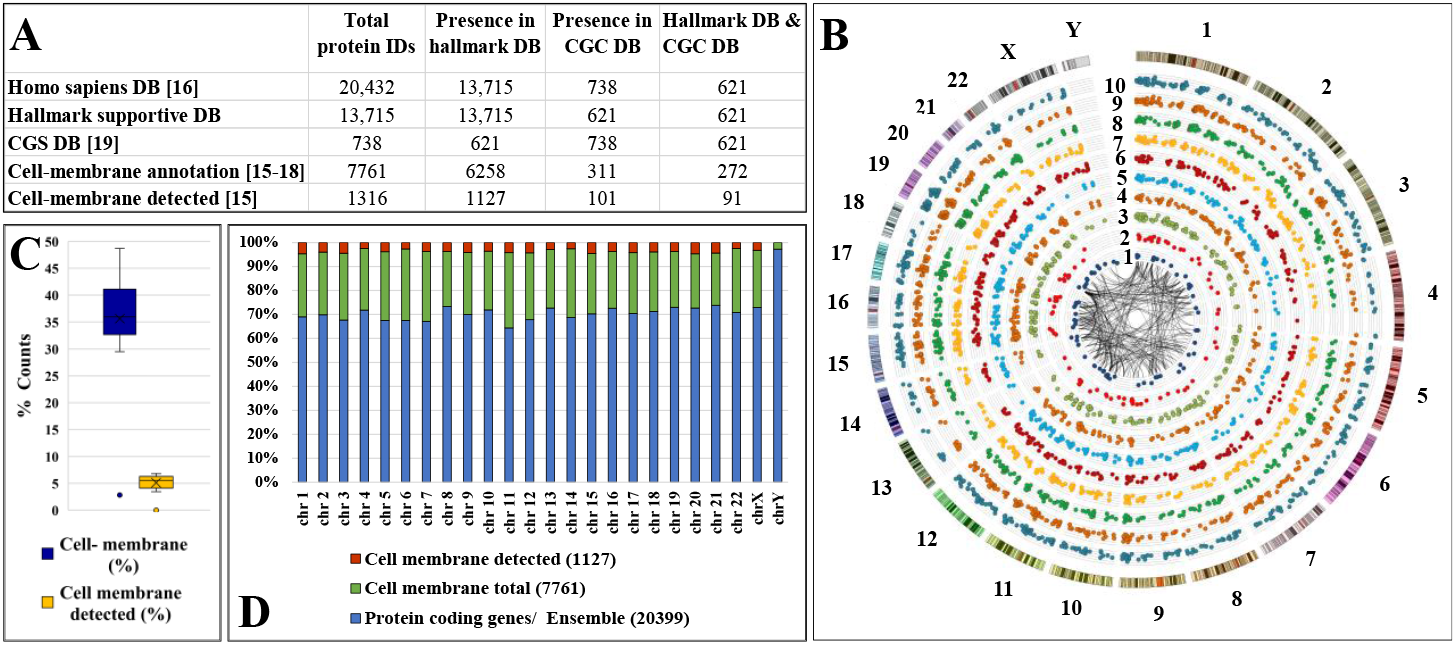
Cell-membrane proteins mapped to cancer hallmarks. (**A**) Protein counts with hallmark association. **(B)** Circos plot of detected SKBR3 cell-membrane proteins mapped to their position in the human genome represented by 23 chromosomes, and categorized into 10 radially distributed cancer hallmarks. Hallmark categories, from inside-out: 1-DNA replication, chromosome organization, and telomeres; 2-glycolysis, gluconeogenesis, and carbohydrate metabolism; 3-Angiogenesis; 4-Inflammatory response; 5-Cell death, apoptosis, senescence, aging; 6-Disease mutation; 7-cell cycle division, growth, and proliferation; 8-Immune system processes; 9-Cell adhesion and motility; 10-Cell communication and signaling. The inner PPI network was constructed with proteins for which the STRING interaction score was ≥0.995. (**C**) Percent counts of cell-membrane proteins encoded on, and detected from, each chromosome. (**D**) Distribution of coded (total and cell-membrane) and detected proteins per chromosome.

To test our hypothesis, a dataset of SKBR3 proteins generated experimentally by three complementary cell-membrane/cell surface enrichment processes [15] was queried for matches to the cancer hallmark DB. Mass spectrometric analysis yielded over 1300 cell-membrane proteins, all identified with high confidence by two unique peptides, of which as many as 1,127 could be matched to multiple hallmarks (**Table 1** and **Supplemental file 1**). A Circos plot with radially distributed hallmarks, in which each cell-membrane protein was mapped to its corresponding gene locus within the 23 human chromosomes, highlights the complex genetic landscape that can lead to cancer through various combinations of genes with altered functional products (**Figure 1B**). The cell-membrane proteins represent ~35-40 % of the encoded proteins by each chromosome, with ~5 % being detectable in SKBR3 when using the described experimental conditions (**Figures 1C&D**). The hallmark-supportive proteins were encoded by the entire genome, excepting chromosome Y which did not have any matches because the SKBR3 cells originate from a female subject (**Supplemental file 1**). Protein abundance was represented by scatter plots depicting the log10(SC), with dot size representing a range spanning from 0 to 5. The non-coding centromeric areas and acrocentric chromosomal p-arms (13, 14, 15, 21, and 22) did not display any products (**Figures 2A&B**), while some chromosomal coding regions appeared to be either under- (4p/4q, 6p, 14q, 17p, Xp, 18q, and 22q) or over-represented (21q) experimentally (**Figures 2C&D**). The 14 proteins encoded by 21q were indicative of positive regulation of ROS metabolic and immune system processes, whereas angiogenic processes emerged more prominently across the entire spectrum as being supported by a range of receptors and adhesion proteins (e.g., THBS1, FGFR1/2, NOTCH1/2, TGFBR1, ERBB, CDC42, CAMs, ephrins, and integrins). The interpretation of such data calls, however, for prudence and further validation, as the SKBR3 cell line is highly aneuploid with numerous structural and numerical chromosomal aberrations (modal chromosome number 84), and the mechanistic impact of chromosomal alterations on protein expression and pathogenic outcomes still lacks a thorough understanding.

**Figure 2.**
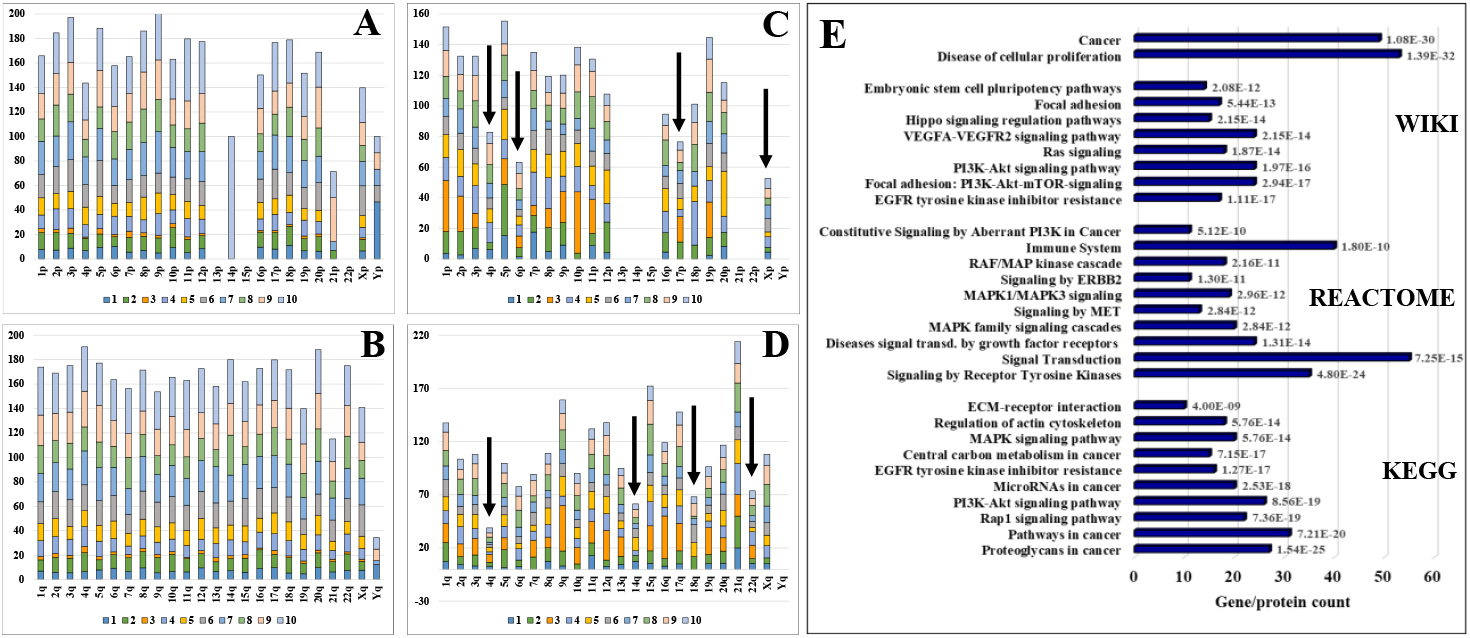
Stacked bar charts representing cancer-supportive human genes and proteins distributed per chromosome arms and per cancer hallmarks; the vertical ordering of hallmarks is following the same numerical trend as in figure 1B (1-chart bottom, 10-chart top). (**A**) and (**B**) represent % cell-membrane protein-coding genes out of total coding genes per chromosome and p/q arms (the summed percentages exceed 100 because of genes with contributions to multiple hallmarks); (**C**) and (**D**) represent % cell-membrane proteins detected in SKBR3 out of total encoded; (**E**) Bar chart of top biological processes and pathways represented by the 116 subset of hallmark proteins (bar chart labels indicate FDRs).

To corroborate the relevance of these cell-membrane proteins to cancer, the COSMIC CGC list was queried against the hallmark database (**Table 1** and **Supplemental file 1**). The CGC comprises evidence-based, manually-curated information related to over 700 cancer-driving genes [19]. The majority of CGC proteins (621) could be mapped to the hallmark database, and encompassed 91 experimentally identified cell-membrane proteins of which 74 were tier 1 (i.e., with documented cancer-relevant activity and evidence of mutations that support oncogenic transformation) and 17 were tier 2 (i.e., with strong indications of cancer-related activity, but lack of sufficient evidence). The detected CGC cell-membrane subset also comprised 44 oncogenes, 20 tumor suppressors, and 8 proteins with overlapping oncogene/tumor suppressor roles. For further evaluation, the list of 91 CGC membrane proteins was supplemented with an additional 25 proteins with documented hallmark roles based on literature reports [1,2,20–24]. Not surprisingly, the top pathways that could be associated with this combined set of 116 representative proteins included cancer-enabling signaling (MAPK, PI3K/AKT, ERBB2, VEGF, HIPPO, EGFR tyrosine kinase inhibitor resistance, stem cell pluripotency) and migration supportive pathways (regulation of actin cytoskeleton, adhesion, ECM-receptor interaction, Rap1) (**Figure 2E**). Multiple small molecule or monoclonal antibody approved cancer drugs that target oncogenes or tumor suppressors from this list have been already described in the DrugBank database (**Figure 3A**).

**Figure 3.**
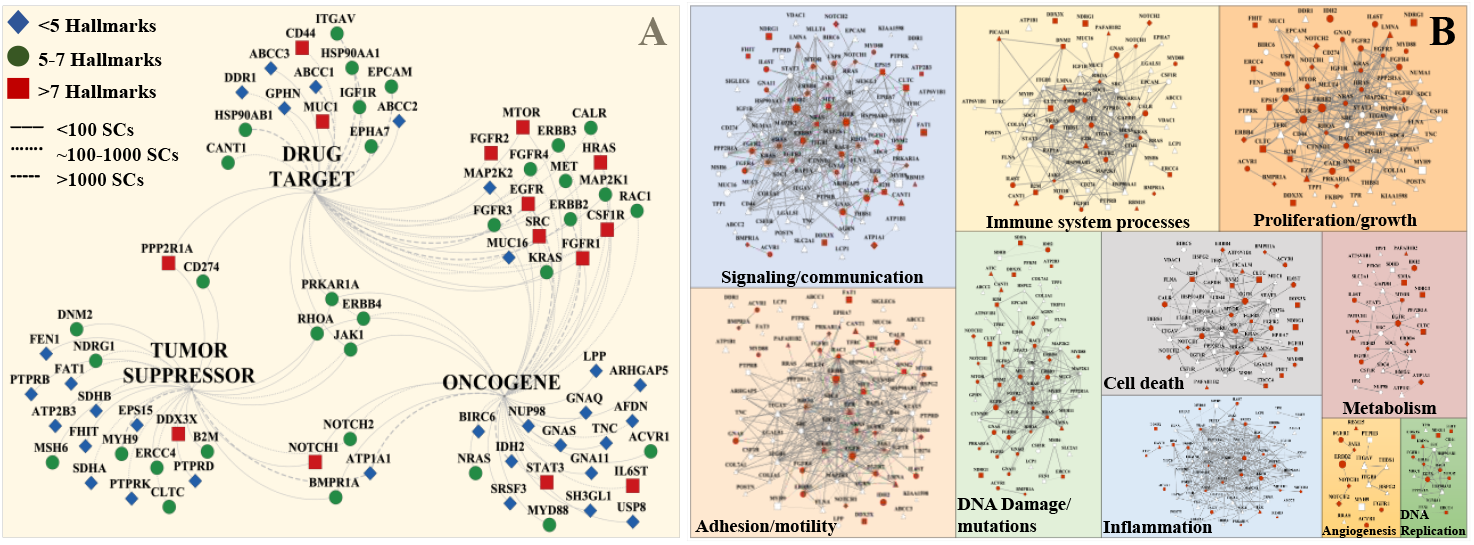
Drug targeting and PPI networks of 116 cell-membrane hallmark proteins. (**A**) Overlapping proteins between SKBR3 cell-membrane oncogenes, tumor suppressors, and drug targets. Top/left legend indicates the symbols of proteins matched to a particular number of hallmarks. Line width indicates the peptide spectral counts matched to each identified protein. (**B**) PPI networks constructed from 116 cell-membrane proteins matched to ten cancer hallmarks. CGC proteins are represented as oncogenes (○), suppressors (•), or oncogenes and suppressors (◇); Red icons indicate whether the protein was annotated as a cancer hallmark protein by CGC; Other hallmark proteins (Δ); Node size is proportional to the protein abundance; Edge thickness reflects the STRING interaction score (≥0.7).

PPI networks constructed based on the detected 116 subset cancer proteins (**Figure 3B**), or based on the whole set of 1,127 proteins (**Supplemental figure 1**), underscored the intimate involvement of the SKBR3 cell-membrane proteome in each of the hallmarks and the progression of cancer. Network analysis of the 116 proteins indicated node degree and betweenness centrality values ranging between 1-32 and 2.88E(-4)-0.2018, respectively, both following the expected power-law distribution (**Figure 4A/**top two panels, and **Supplemental file 1**). Circular layout representations of degree and betweenness were created for the proteins that had interacting partners (81 out of 116), and revealed that the targets of the existing cancer drugs are proteins with rather high degree than high betweenness (**Figure 4B** and **C**). With only 50 proteins displaying a betweenness value >0, a correlation between betweenness and degree was observable mainly for degree values >10 (**Figure 4A**/bottom panel). Certain proteins, such as ERBB4, were characterized by a high degree (=7) but low betweenness (=0). Correlation between protein abundance and propensity for targeting was, however, not evident.

**Figure 4.**
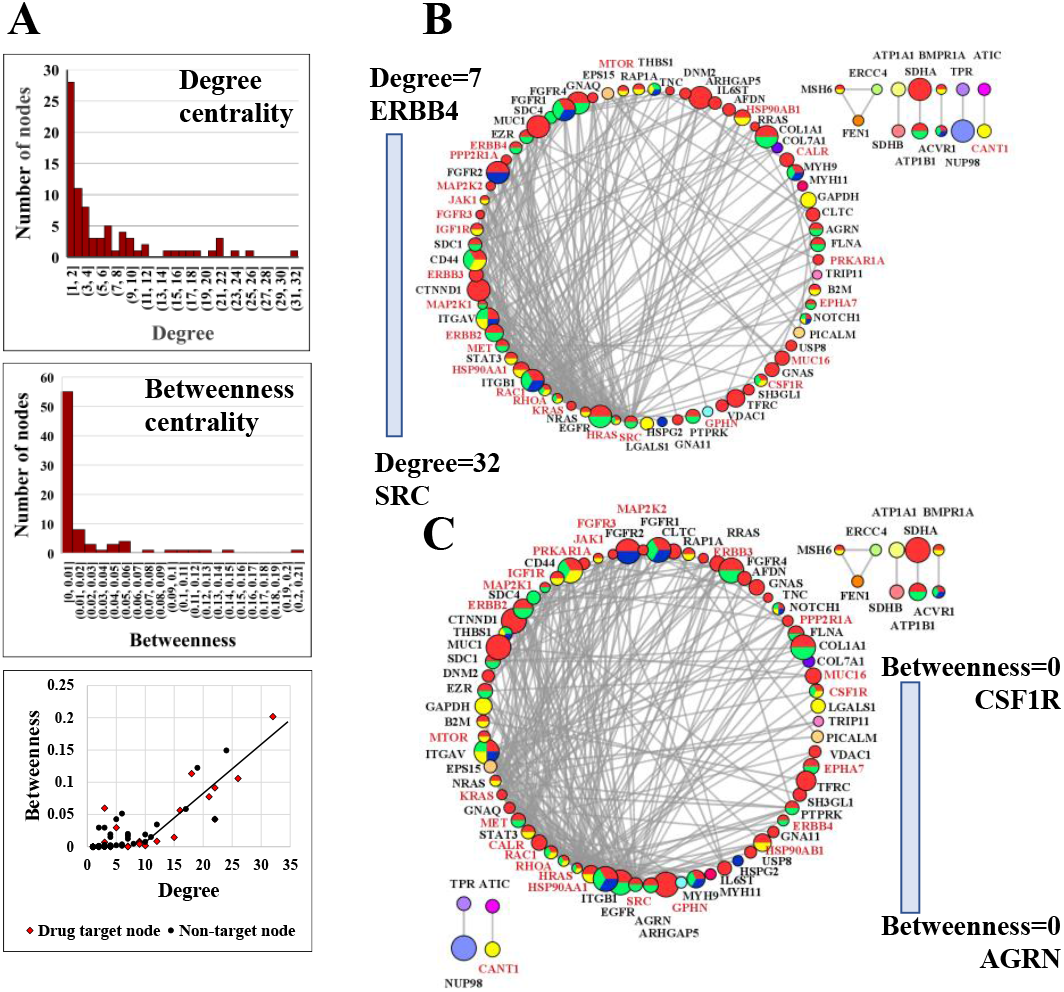
Network analysis of PPI networks created from 116 SKBR3 cell-membrane proteins (STRING interaction score ≥0.9). (**A**) Distribution and correlation of degree and betweenness centrality measures. (**B**) Degree-based circular layout PPI network (degree=1-32). (**C**) Betweenness-based circular layout PPI network (betweenness=2.88E(-4)-0.2018). The circular distribution of attributes is presented in a clockwise fashion, from high (left) to low (right). Node size is proportional to the protein abundance and represented as LOG10(SC) over a range of 0.699-4.217. Node color coding: red-signaling, yellow-immune response, green-locomotion, blue-angiogenesis. Gene names that represent drug targets are shown in red. Network statistics: nodes 116 (only 81 nodes with degree ≥1 are shown), edges 268, avg # neighbors 7.5, diameter 6, radius 3, clustering coefficient 0.44, density 0.11.

## Discussion

All major cell-membrane protein categories, i.e., with receptor/catalytic activity, adhesion/junction, transport, and CD classification, were represented in every cancer hallmark. The hallmarks of cell communication/signaling, adhesion/motility, immune response and cell cycle/growth accrued the largest number of protein hits, reflecting their considerable impact on cancer development and dissemination (**Table 1**). Some of the highest abundance membrane proteins were part of the very same categories, and included receptors, adhesion proteins, solute carriers, and transporters, as well as additional ephrins, nectins, integrins and essentially all detected CDs. A few specific examples, that also comprise hallmark proteins from the classic literature (highlighted in bold), included members with multiple roles in receptor mediated signaling (**ERBB2/CD340**, **EGFR**, TFRC/CD71, PTPRF, ITGAV/CD51, ITGB1, BCAM, PVR, SCARB1, SUSD2), adhesion (ITGAV, ITGB1/CD29, BCAM, **EPCAM/CD326**, PVR, SCARB1, MUC16, PVR, PTPRF, **EGFR**, **ATP1B1**, SCARB1), immune response (SUSD2, PVR, SCARB1, ITGAV, **EGFR**, TFRC/CD71, CD44), and transport (ABCC1, ATP1A1, **ATP1B1**).

The PPI networks from **Figure 3B** exposed highly interconnected protein clusters with several members playing roles in multiple hallmark categories (e.g., EGFR, ERBB2, FGFRs, CD44, SRC, STAT3). The emerging information can be used to identify novel regulatory or effector proteins, targets for knockout experiments, components of protein complexes, or to assign functionality to yet uncharacterized proteins. Through PPI networks, attention can be expanded to a vast set of membrane proteins that when targeted in a combinatorial or sequential fashion can impair cancer proliferation and dissemination not only through well-established mechanisms such as RAS/ERK or PI3K/AKT signaling, but also via processes implicated in angiogenesis, migration, cell death, metabolic reprogramming, inflammation, or remodeling of the tumor microenvironment. The ability to identify by MS aberrant protein isoforms or posttranslational modification patterns, e.g., glycosylation in the case of cancer [29], can further strengthen the efficacy of this approach. Pathways that confer intrinsic or acquired drug resistance and that often work via cross-talk or transactivation, or that lead to cancer stem cell mediated resistance, have been of particular interest to drug developers. Informed selection of drug targets, rather than based on empirical screening, will better suppress redundant or compensatory signaling pathways, prevent tumor-induced vascular remodeling, disable the ECM supportive environment, overturn immune evasion capabilities, and impair metastatic potential.

The SKBR3 cell-membrane landscape of oncogenes, tumor suppressors and existing drug targets revealed one such example of a group of key receptors (EGFR, ERBB, FGFR, IGFR, MET, MTOR, MAPK/RAS proteins), CD antigens (CD274, CD44), adhesion proteins (EPCAM, MUC1), and ABC transporters (ABCC1/2/3) that have been already explored for effective targeting of various cancers (**Figure 3A**). Such complex protein signatures and network architectures enable the targeting of broad panels of interconnected RTKs or CDs, and can uncover vulnerabilities that may trigger cascading failures from either within the established (**Figure 3B**) or the extended networks (**Supplemental figure 1**). This, in turn, can enable combating widespread strategies used by cancer cells to resist the action of drugs, such as MET amplification or PI3K/AKT activation after targeted EGFR inhibition, PI3K/AKT compensation after mTOR inhibition, or MAPK/STAT3 cross-talk [14]. Targeting orthogonal pathways by using combinations of signal transduction or angiogenesis inhibitors (e.g., targeting Tyr kinase oncogenes), regulators of apoptosis (e.g., MUC1/16, a negative regulator of intrinsic apoptotic pathways), and inhibitors of drug efflux proteins (e.g., ABCC1 or multidrug resistance associated protein-MRP1), can be further explored to selectively attack or eliminate the cancer cells.

Network pharmacology, the topology of such networks, and the centrality of drug targets in these networks has been extensively researched to help identify and validate novel proteins amenable to therapeutic targeting [30–34]. In silico prediction of drug targets has been explored, for example, by using a multilayered interactome analysis coupled with a “guilt-by-association” approach [30]. While the known cancer drug targets in the circular diagrams from **Figures 4B** and **4C** were distributed along the entire range of degree and betweenness centrality values, the targets correlated better with high degree than high betweenness (note the drug target gene names shown in red). Most cell-membrane or cell-membrane associated drug targets (16 out of 24) included proteins with a degree ranging from 32 (SRC) to 7 (ERBB4), in the upper third of the degree distribution scale (**Figure 4B**). This highly interconnected set of 16 drug targets is involved mainly in intracellular signaling and positive regulation of kinase activity, the most represented KEGG pathways being EGFR Tyr kinase inhibitor resistance (11 proteins), ERBB (10 proteins), and PI3K-AKT signaling (13 proteins). In contrast, many targets fell in a region of the circular diagram where betweenness was minimal (**Figure 4C**). Proteins from the lower third of the betweenness scale did not form PPI networks. This result supports previous predictions that pinpointed that node degree in non-directed PPI networks is a better predictor of essentiality than betweenness, because there is no information flow through the nodes such as in the case of signal transduction networks [34]. Using centrality metrics in the context of a more limited but more fertile class of proteins, such as encompassed by the cell-membrane proteome, represents a promising alternative for identifying new drug targets. For example, several proteins for which drug antagonists have not yet been developed or approved, and that line up in the circular diagram with degree=5-7, have been just recently suggested for consideration as molecular targets. Among these, EZR has been proposed as a prognostic marker and target in acute myeloid leukemia [35], MUC1 for the development of anti-MUC1 antibodies for targeted therapy of GI cancers and development of anti-cancer vaccines [36], SDC4 for hepatocellular carcinoma [37], and FGFR4-also associated with resistance to anti-tumor therapy-for breast cancer [38]. Moreover, as aberrations in G-protein activating subunits have been shown to act as driver mutations implicated in multiple cancers [39], the use of siRNA to target and downregulate mutated GNAQ in several cancers, in particular melanoma, has been also explored [40]. These novel trends underscore the power of orthogonal discovery efforts that can deliver valuable candidates for supporting the development of synergistic cancer therapeutic approaches that overcome the simplistic “one disease/one target/one drug” paradigm.

As a complementary undertaking, tumor-stroma interactions that support cancer progression and metastasis can be interrupted by disabling paracrine (FGF, Wnt, Hedgehog, TGFβ, NOTCH) and autocrine signaling sustained by cell-membrane receptors and proteases. Therapies aimed at modulating the tumor-immune cell dialogue, by using for example check point inhibitors such as the anti-programmed cell death ligand 1 (PD-L1 or CD274) or anti-PD1, with PD-L1 also detected among the SKBR3 membrane proteins, are expected to find a fertile ground in the rich backdrop of cell-membrane antigens. Such immunotherapies can be used alone or jointly with chemo- or targeted therapies.

## Conclusions

In summary, mapping the SKBR3 cell-membrane proteome to cancer hallmark-supportive processes exposed a complex scenery of critical players that promote and sustain the development and metastatic propensity of these cells. The cell-membrane harbors an abundant supply of putative targets for the development of modern cancer therapies, future work being needed to refine the list of viable candidates. The analysis of even one single cell line was informative enough to uncover a large number of existing oncogene and tumor suppressor drug targets, and through protein-protein interactions and network analysis to reveal that network centrality measures are useful indicators for the selection of new targets. With the advance of high-throughput MS instrumentation, the ability to quickly generate tumor cell-membrane profiles will create the necessary resource for exploring complex and aberrant protein expression signatures, studying additive or synergistic effects of drug combinations that target hallmark proteins, understanding the evolution of (multi)drug resistance, evaluating cancer risk and prognostic factors, and facilitating the selection of patient-tailored drug targets for achieving superior therapeutic outcomes in the various stages of cancer development.

## Acknowledgment

This work was supported by an award from the National Institute of General Medical Sciences to IML (R01 Grant No. GM121920).

## Author contributions

AK processed and analyzed the mass spectrometry data, reviewed and queried the literature for cancer hallmark proteins, and constructed the PPI networks. JRSH analyzed the hallmark protein dataset and generated the Circos plot. IML conceptualized the work, built the cancer hallmark protein database, performed the network analysis, and wrote the manuscript. All authors reviewed and approved the final version of the manuscript.

## Correspondence

Address correspondence and material requests to Dr. Iulia M. Lazar.

## Data availability

Supplemental file 1 comprising the protein IDs associated with the cell-membrane cancer hallmark processes and the detected proteins and degree/betweenness centrality measurements, as well as Supplemental figure 1 providing the extended PPI networks, are available through the journal website. The mass spectrometry raw files are available through the ProteomeXchange Consortium via the PRIDE partner repository with the following dataset identifiers: PXD028976, PXD028977, and PXD028978.

