## Supplemental figure 1 for "Mapping the Cell-Membrane Proteome to the Cancer Hallmarks"

### Slide 1
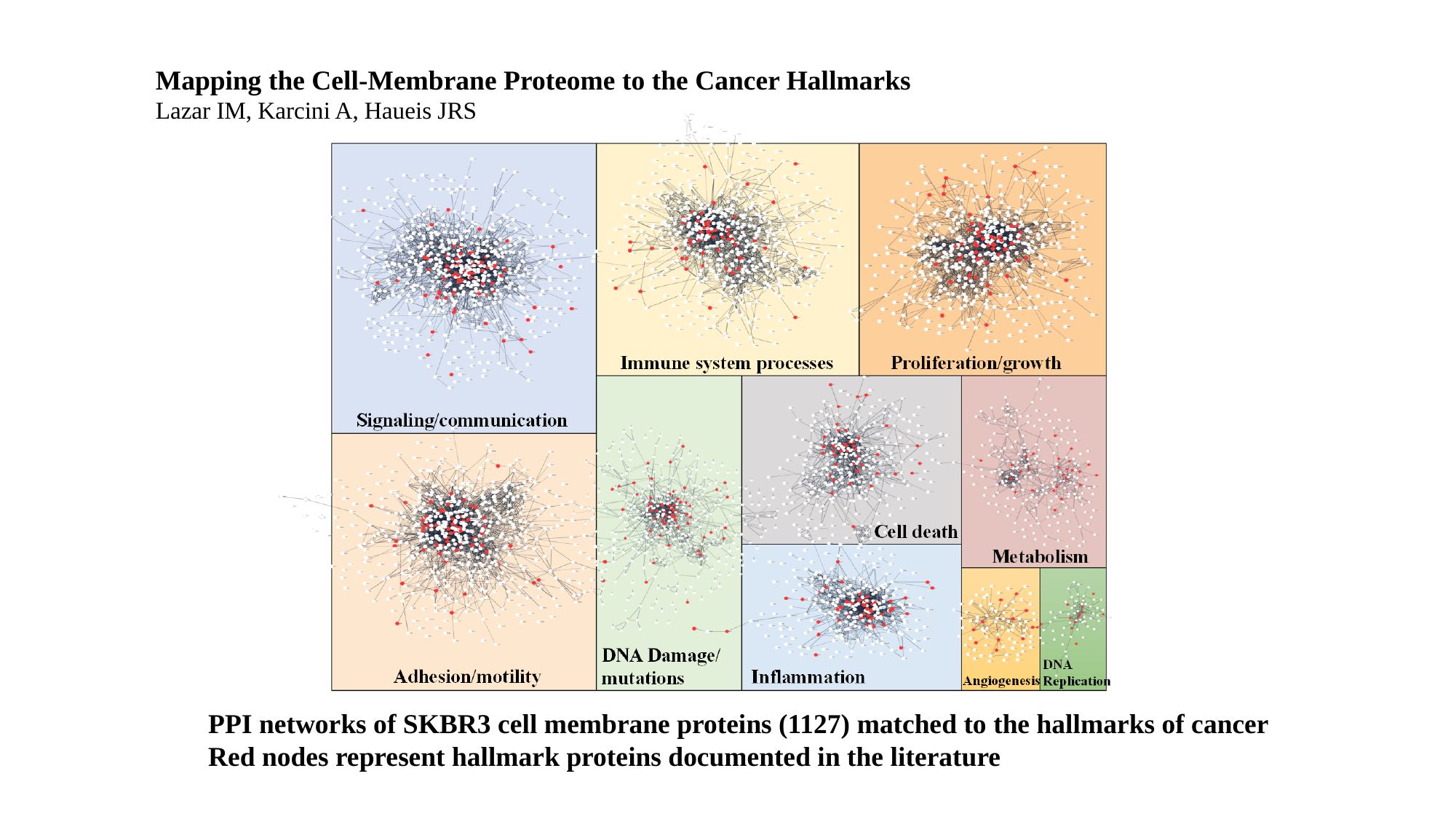

Mapping the Cell-Membrane Proteome to the Cancer Hallmarks
Lazar IM, Karcini A, Haueis JRS
PPI networks of SKBR3 cell membrane proteins (1127) matched to the hallmarks of cancer
Red nodes represent hallmark proteins documented in the literature
